# Evolutionary transition in accessible chromatin landscapes during vertebrate embryogenesis

**DOI:** 10.1101/481309

**Authors:** Masahiro Uesaka, Shigeru Kuratani, Hiroyuki Takeda, Naoki Irie

## Abstract

The relationship between development and evolution is a central topic in evolutionary biology^1,2^. Recent transcriptome-based studies support the developmental hourglass model, which predicts that the animal embryogenetic program is most strongly conserved at mid-embryonic stages^3-9^. This model does not necessarily contradict the classical hypothesis^10,11^ that animal development recapitulates its evolutionary history after the mid-embryonic stages^2,12^. However, to date there is no molecular evidence supporting the hypothesis that gene-expression profiles that are more evolutionarily derived appear sequentially in late development. Here, by estimating activated genomic regions and their evolutionary origins, we show that the recapitulative pattern appears during late embryonic stages. We made a genome-wide assessment of accessible chromatin regions throughout embryogenesis in three vertebrate species (mouse, chicken, and medaka) and determined the phylogenetic range at which these regions were shared. In all three species, sequential activation of putative regulatory regions that were more derived occurred later in embryogenesis, whereas ancestral ones tended to be activated early. Our results clarify the chronologic changes in accessible chromatin landscapes and reveal a phylogenetic hierarchy in the evolutionary origins of putative regulatory regions that parallels developmental stages of activation. This relationship may explain, at least in part, the background for morphological observations of recapitulative events during embryogenesis.

---

The idea of a parallel relationship between development and evolution traces back to the work of von Baer^10^ and Haeckel^11^ in the 19th century. Haeckel’s model—recapitulation—suggests that animal development proceeds along a phylogenetic evolutionary pathway, sequentially developing from features that are more ancestral to those that are more derived. However, recent transcriptome-based studies supporting the hourglass model have not shown such a recapitulative pattern in the early-to-late embryogenesis of vertebrates^3,4,6,8,9^. Rather, after conserved mid-embryonic stages, or the phylotypic period^13,14^ (around E9.0 in mouse^6,9^, around HH16 in chicken^6,8,9^ and around stage 24 in medaka; Extended Data Figure 1 and Supplementary Note 1), vertebrate embryogenesis might still mirror the process of phylogenetic trajectory, recapitulating its evolutionary history^2,12^. The development of some morphological features that appear after the pharyngula stage, such as loss of limbs in snakes, turtle shell formation^15^ and jaw development in needlefishes^16^, could be explained by such a scheme.

**Figure 1.**
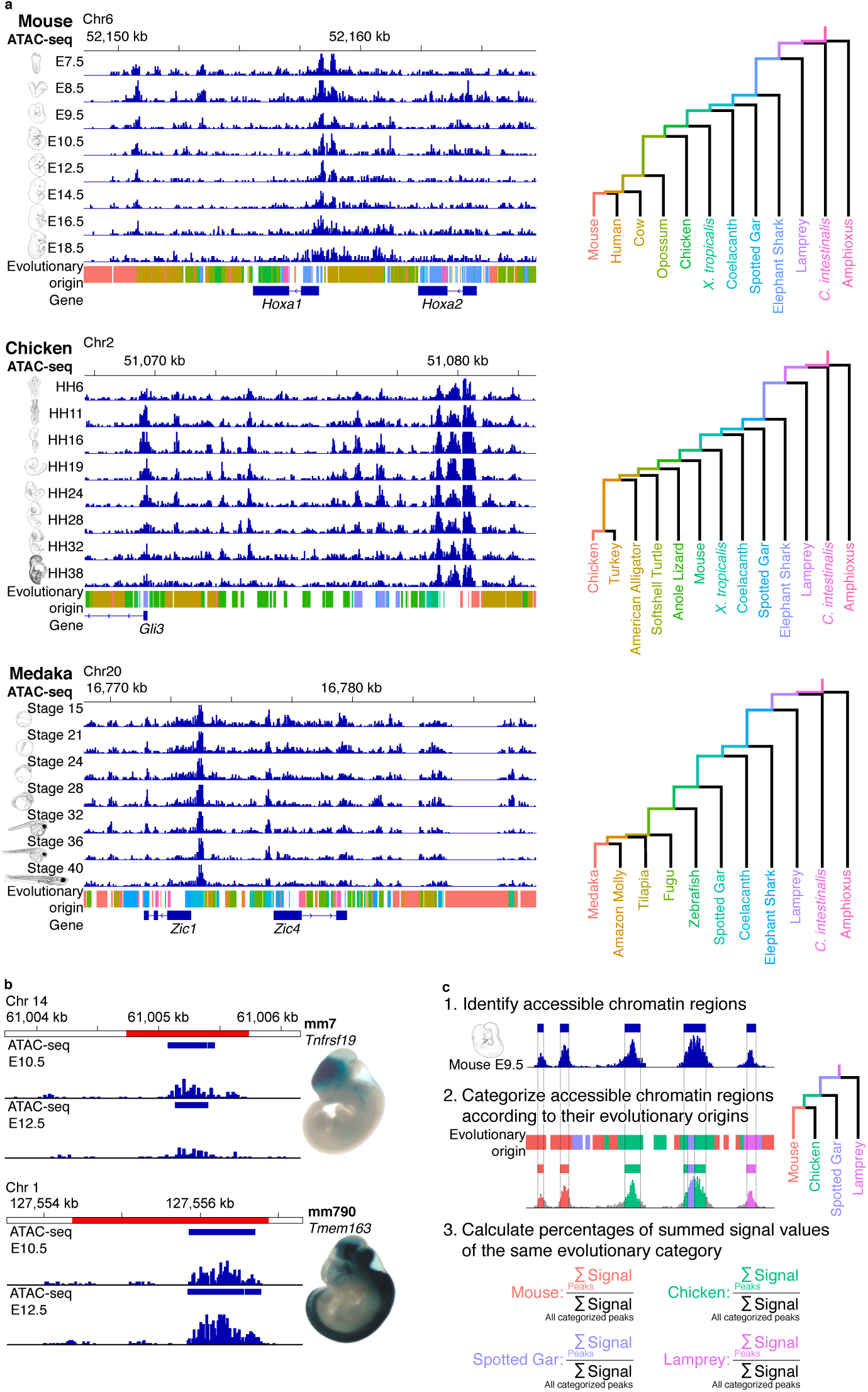
Strategy for assessing accessible chromatin landscapes in vertebrate embryos. **a.** Genome browser views showing whole-embryo ATAC-seq signals in mouse, chicken and medaka embryos. ATAC- seq signals are presented as means of three biological replicates. Colours below the ATAC-seq signals represent the estimated evolutionary origins of genomic regions that correspond to the tracks of the evolutionary trajectories, which are shown as a tree on the right. The evolutionary origin of each region was estimated for the largest monophyletic group sharing the regions with similar sequences. Uncoloured regions indicate regions that were lost in some lineages or species and that were removed for further analyses (see Methods). Phylogenetic relationships and split times were adopted from the TimeTree database^29^. **b.** Identified accessible chromatin regions (blue boxes) overlapping with annotated enhancers (red regions) acquired from the VISTA Enhancer Database^30^. Whole-embryo ATAC-seq signal intensities were correlated with *in vivo* enhancer activity (*LacZ* reporter assay) in E11.5 transgenic mice (see also Extended Data Figure 4). Images of embryos are from the VISTA Enhancer Database^30^. For each annotated enhancer, the VISTA enhancer ID and flanking gene are shown above the embryo image. **c.** Schematic diagram showing the steps for estimating the total chromatin accessibility for each evolutionary origin. Step 1: Accessible chromatin regions were identified according to ATAC-seq signal intensity. Step 2: Identified regions were categorized according to their estimated evolutionary origins. Step 3: For each developmental stage, the percentage corresponding to the summed signal value of each evolutionary category divided by the total signal for all of the evolutionary categories was calculated.

Although the expression profiles of protein-coding genes do not match the recapitulative pattern of vertebrate embryos^3,4,6,8,9^, changes in the epigenetic states of gene regulatory regions might show such a pattern. The majority of genetic changes associated with the emergence of major vertebrate groups are concentrated in potential regulatory regions, not in protein-coding regions^17,18^, and repeated recruitment of the same protein-coding genes at different developmental stages^9^ would obscure any recapitulative pattern, if present. The repertoire of protein-coding genes exhibits far fewer changes than putative regulatory regions in the species we analysed (Extended Data Figure 2), indicating that examining their epigenetic states during development may provide greater resolution for detecting recapitulation. Here, we hypothesize that the evolutionary origins of the activated regulatory regions show sequential changes along the chronological axis of development, which we tested by focusing on epigenetic states in the embryos of three vertebrate species.

**Figure 2.**
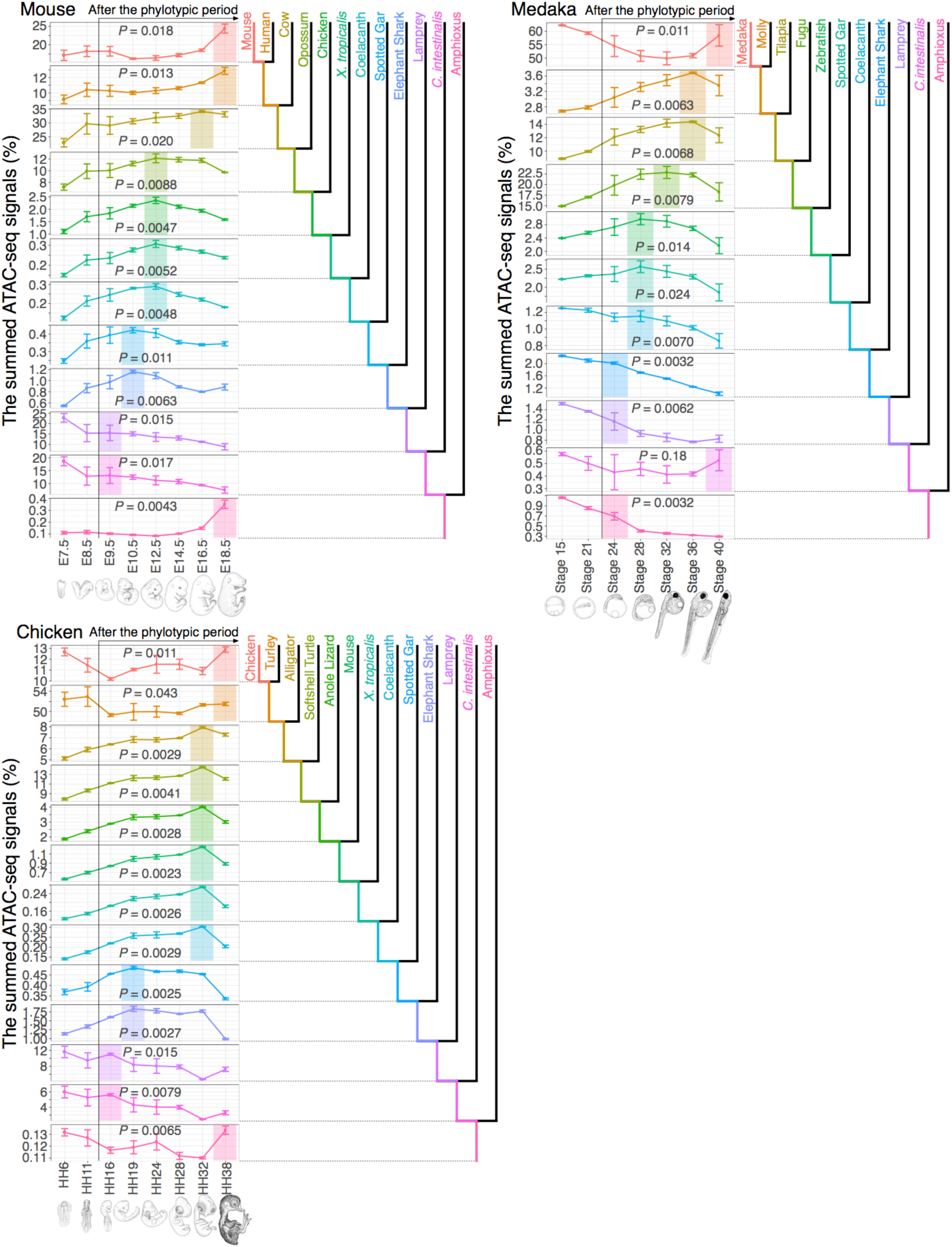
Transition of signal peaks in evolutionarily categorized chromatin accessibilities during vertebrate embryogenesis. For each developmental stage in three vertebrate species (mouse, chicken, and medaka), the percentages corresponding to the summed signal intensity for each evolutionary category divided by the total signal intensity for all categories are shown. The colour of each category indicates the period of evolutionary origin of the region, as represented in the phylogenetic trees adjacent to the graphs. In each graph, the developmental stages with the highest value after the potential phylotypic period are highlighted in the corresponding colour. Error bars indicate the standard deviation of three biological replicates of each developmental stage. The *P* value shown in each graph was calculated by using the Kruskal–Wallis rank sum test. Sample sizes and detailed information regarding the statistical tests (test statistic, degrees of freedom and *P* value) are in Supplementary Table 3.

We collected mouse (*Mus musculus*), chicken (*Gallus gallus*) and medaka (*Oryzias latipes*) embryos for the stages covering the conserved stage and later developmental stages (E7.5–E18.5 in mouse, HH6– HH38 in chicken and stages 15–40 in medaka; Supplementary Table 1). Active regulatory regions were then estimated by using the assay for transposase-accessible chromatin using sequencing (ATAC-seq^19^; Figure 1a), as accessible chromatin marks active regulatory regions, including enhancers, silencers, and promoters^20,21^.

We identified more than 100,000 accessible chromatin regions as putative regulatory regions for each developmental stage (Figure 1a and Extended Data Figure 3a). The chromatin accessibility of these regions was robustly reproduced across the biological replicate samples (Extended Data Figure 3b). We confirmed that several representative accessible chromatin regions overlap with the annotated enhancers (Figure 1b and Extended Data Figure 4). We also evaluated the signal intensity of whole-embryonic ATAC-seq at each accessible chromatin region, as we predicted that this would reflect the number of cells in which the examined region was accessible. Consistently, ATAC-seq signal intensity tended to be stronger for enhancers driving expression in larger populations of cells (Figure 1b and Extended Data Figure 4).

We next traced the evolutionary origins of these accessible chromatin regions by using pairwise whole-genome alignments to the genomes of different chordate species (Supplementary Table 2). For each accessible chromatin region, the origin of the largest monophyletic group that shared the regions with similar sequences was defined as its evolutionary origin (Figure 1c). Single accessible chromatin regions consisting of at least two stretches of sequences with different evolutionary origins were subdivided and evaluated as separate regions (Figure 1c).

Among the identified accessible chromatin regions with evolutionary origins at each developmental stage (more than 250,000 regions for each stage), fewer than 5% were older than the vertebrate–tunicate split (Extended Data Figure 2). In contrast, the same analysis of the protein-coding sequences showed that more than two-thirds of these regions emerged before that split (Extended Data Figure 2), implying that the regulatory region repertoire is much more diverse than that of protein-coding genes, as previously reported^17,18^.

To estimate the relative regulatory activity of each evolutionary origin, we summed the ATAC-seq signal intensities for accessible chromatin regions sharing the same evolutionary origin and calculated the percentage of that sum relative to the total signal intensities for all accessible chromatin regions with evolutionary origins (Figure 1c). For embryogenetic stages later than the phylotypic period^6,8,9^ (stages from E9.5 in mouse, HH16 in chicken and stage 24 in medaka), the more recently acquired accessible chromatin regions consistently tended to show signal percentage peaks at later developmental stages in all three species examined (Figure 2). We obtained essentially similar results for the same analyses using different alignment parameters (Extended Data Figures 5 and 6), a different method for estimating the evolutionary origins of regions (Extended Data Figure 7), and a different set of genomes that covered a wider phylogenetic range (Extended Data Figures 8 and 9), which suggests the overall tendency is robust against differences in analytical parameters (Supplementary Note 2). These results support the idea that transitional patterns of chromatin accessibility parallel the evolutionary pathway. In contrast, as inferred from previous reports^8,9^, analysis of whole embryonic gene expression profiles did not reveal a similar tendency (Extended Data Figure 10).

Before the phylotypic period, however, this recapitulative tendency in chromatin accessibility was not apparent in chicken or medaka (HH6 to HH16 in chicken and stages 15 to 24 in medaka; Figure 2 and Extended Data Figures 5–9); these results are consistent with the idea^2,12^ that the recapitulative pattern likely would not be apparent in early embryogenesis. In mouse, however, this pattern was observed in embryogenesis, even from E7.5 (Figure 2 and Extended Data Figures 5–9). Although the first stage to show the recapitulative pattern may vary by species, it also is possible that the accessible chromatin landscape of mouse embryos earlier than E7.5 does not recapitulate phylogeny, as those of chicken and medaka embryos do, given that the E7.5 stage in mouse shows high conservation at the transcriptome level and therefore could be included in the phylotypic period^9^. Moreover, in all three species, recently acquired regions showed higher levels of activity at early developmental stages than at mid-embryonic stages (Figure 2 and Extended Data Figures 5–9). These results are in accord with previous reports, which show that sequences of active regulatory regions are more evolutionarily conserved in cell lines and in organs derived from mid-stage embryos than in those from early-stage samples of humans and mouse, respectively^22,23^. Our findings suggest that putative recapitulation can be observed in regulatory activity after the phylotypic period.

Despite this putative recapitulation in accessible chromatin landscapes, results for evolutionarily old origins differed only slightly by species. Considering the observed evolutionary range over the recapitulative tendency, for mouse and chicken, the transitional pattern of chromatin accessibility of regions newer than the split from the amphioxus exhibited recapitulation. In contrast, during medaka embryogenesis, this pattern consistently emerged for those acquired after the cyclostome–gnathostome split (Figure 2 and Extended Data Figures 5–9; Supplementary Note 2.4). This inconsistency between medaka and other species could be due in part to genome-level changes during medaka evolution (Supplementary Note 3). However, for accessible chromatin regions newer than the cyclostome–gnathostome split, this recapitulative tendency was observed in all of the species examined, consistently across different analytical parameters (Figure 2 and Extended Data Figures 5–9). These findings suggest that, in all of the species examined, late embryogenesis shares the same recapitulative tendency of chromatin accessibility, at least for regions that emerged more recently than the cyclostome– gnathostome split.

Our findings show that the epigenetic states of vertebrate genomes sequentially follow a pattern that proceeds from ancestral to derived states during development. This orchestration could potentially underlie key developmental factors behind some phenomena recognized as recapitulation by comparative embryologists. At the morphological level, our results are consistent with previous studies^2,12,15,16^ that showed recapitulation-like development of several morphological features acquired during vertebrate evolution. Puzzlingly, no similar tendency has been detected at the level of the transcriptome. This may be attributable to a focus on changes in the protein-coding gene repertoire during development, which would be insufficient to examine the evolutionary trajectories of vertebrate embryogenesis.

If this recapitulative tendency holds true for vertebrate embryogenesis, it may indicate that, after the acquisition of new regulatory elements in the ancestral genome, the activation of such elements at earlier developmental stages would lead to lethal or less adaptive phenotypes more often than later activation. This is consistent with similar rationales suggested in several other authors’ hypotheses^24-27^ and the results of a simulation-based study^28^; i.e., that developmental processes at earlier stages establish the foundations for later ones, and that the existence of newly acquired processes presupposes preceding ones on which they depended at earlier stages. This, of course, may not always be the case, as evolutionarily recently acquired regions were activated at earlier stages than the middle period of development. It would be of interest to examine how these changes have become integrated into stages of embryogenesis earlier than the conserved phylotypic period. Further studies are needed to resolve the mechanisms underlying the recapitulative tendency we describe here and its relationship with the developmental hourglass model. To this end, molecular and genomic analyses are needed to connect evolutionary changes in gene regulation and their effects on embryonic divergence in the context of modern evolutionary biology.

## Supplementary Notes

This file contains Supplementary Notes 1–3 and additional references.

## Supplementary Tables

This file contains Supplementary Tables 1–11.

### Acknowledgements

This work was supported in part by JSPS KAKENHI Grant Numbers JP15J06414 (Grant-in-Aid for JSPS Research Fellow) and JP18K14711 (Early-Career Scientists) to M.U.; JSPS KAKENHI Grant Numbers JP15H05603 (Grant-in-Aid for Young Scientists (A)) and JP17H06387 (Grant-in-Aid for Scientific Research on Innovative Areas) and the Platform for Dynamic Approaches to Living System from the Ministry of Education, Culture, Sports, Science and Technology, Japan to N.I.; and JSPS KAKENHI Grant Number JP17H06385 (Grant-in-Aid for Scientific Research on Innovative Areas) and a Naito Grant for the Promotion of Focused Research (The Naito Foundation) to S.K. We thank Ryohei Nakamura, Masahiko Kumagai and Yui Uchida for useful discussions and help in performing ATAC-seq. We thank Tatsuya Hirasawa, Juan Pascual-Anaya, Takahiro Kohsokabe, Rie Kusakabe and Douglas Sipp for critical reading of the manuscript. Computations were partially performed on the NIG supercomputer at ROIS National Institute of Genetics and on the Bioinformatics Analysis Environment Service on RIKEN Cloud at RIKEN Advanced Center for Computing and Communication.

## Author contributions

M.U. and N.I. conceived and designed the study with important help from H.T. and S.K. M.U. collected the samples with extensive help from H.T., conducted the experiments and analysed the data. All authors interpreted the data. M.U., N.I. and S.K. wrote the paper. All authors approved the final version of the manuscript.

## Author Information

All data have been deposited in the DNA Data Bank of Japan with the accession number DRA006971. The authors declare no competing interests. Readers are welcome to comment on the online version of the paper. Correspondence and requests for materials should be addressed to U.M.

## Methods

### Data reporting

No statistical methods were used to predetermine sample size. The investigators were not blinded to allocation during experiments or during outcome assessment.

### Animal care and use

Experimental procedures and animal care were conducted in accordance with the guidelines approved by the Animal Experiment Committee of the University of Tokyo (Animal_plan_26-3). All efforts were made to minimize animal pain and distress. Individual embryos were selected blindly from a wild-type population.

### Embryo collection

#### Mus musculus

All embryos were collected from C57BL/6J mice (CLEA Japan) and staged according to standard morphological information on normal mouse developmental stages^31^. After the amniotic membranes had been removed from the staged embryos, we pooled at least two embryos from different pregnant mice to prepare each biological replicate (Supplementary Table 1).

#### Gallus gallus

NERA-strain fertilised chicken eggs were purchased from a local farmer in Japan (Shiroyama- keien, Kanagawa, Japan). Fertilised chicken eggs were incubated at 38 °C in a humidified incubator and morphologically staged according to the Hamburger– Hamilton system^32^. After the amniotic membranes had been removed from staged individual embryos, we pooled at least two embryos to prepare biological replicates of each developmental stage (Supplementary Table 1).

#### Oryzias latipes

Mature adults of the d-rR strain were maintained under standard conditions (10:14-h dark:light cycle; 26–28 °C) and mated to obtain fertilised eggs. Fertilised eggs were incubated at 24–26 °C, and individual embryos were morphologically staged, as described by Iwamatsu^33^. Biological replicates comprised pooled embryos from different pairs of parents (Supplementary Table 1).

### Preparation and sequencing of ATAC-seq library

For each biological replicate, ATAC-seq was performed as previously described^34,35^, with some modifications. In brief, embryos were minced by using razor blade as required, placed in homogenization buffer (25 mM D-sucrose, 20 mM tricine [pH 7.8], 15 mM NaCl, 60 mM KCl, 2 mM MgCl2, 0.5 mM spermidine, and cOmplete Protease Inhibitor Cocktail Tablet [Roche]), and homogenized in an ice-cold Dounce tissue grinder with a loose-fitting pestle, according to the methods of Yue *et al*.^36^. For further dissociation, each homogenized sample was forced through a 21-gauge needle by using a syringe and then filtered through a 100-µm Filcon filter (As One Corporation). Cells were harvested by centrifugation at 500 × *g* for 5 min at 4 °C and then resuspended in 500 µL of cold sucrose buffer (250 mM D-sucrose, 10 mM Tris–HCl [pH 7.5], 1 mM MgCl_2_, and cOmplete Protease Inhibitor Cocktail Tablet [Roche]). In total, 500,000 cells or 50,000 cells was centrifuged at 500 × *g* for 5 min at 4 °C for each sample, depending on the developmental stage (Supplementary Table 1). Cells were re-suspended in 50 μL cold lysis buffer (10 mM Tris–HCl [pH 7.4], 10 mM NaCl, 3 mM MgCl_2_, 0.1% v/v Igepal CA-630 [Sigma– Aldrich]) and incubated on ice for 5 min. After centrifugation of the cells at 500 × *g* for 10 min at 4 °C, the supernatant was discarded. Tagmentation reactions were performed at 37 °C for 30 min by using a Nextera Sample Preparation Kit (Illumina). Tagmentated DNA was purified by using a DNA concentrator kit (Zymo Research) and size-selected (<500 bp) by using AMPure XP beads (Beckman Coulter). Next, to enrich for small DNA fragments, two sequential PCR amplifications were performed, as described previously^34^. The amplified DNA was purified by using a DNA concentrator kit (Zymo Research), and the quality of the purified library was assessed by using a 2100 Bioanalyzer (Agilent Technologies) and a High Sensitivity DNA analysis kit (Agilent Technologies) to confirm a periodic pattern in the size of the amplified DNA. After size selection (100–300 bp) by using AMPure XP beads (Beckman Coulter), the libraries were sequenced as paired-end 50-bp reads by using the HiSeq 1500 platform (Illumina) according to the manufacturer’s instructions.

### Alignment of ATAC-seq reads

Adaptor trimming and quality filtering of raw paired-end reads were performed by using Trimmomatic (version 0.36)^37^ with the following parameters: ‘ILLUMINACLIP: adaptor.fa:2:30:10, LEADING: 20, TRAILING: 20, SLIDINGWINDOW: 4:15, MINLEN: 36’. The filtered reads were aligned to species-specific reference genomes (GRCm38 for *Mus musculus*^38^, Gallus_gallus-5.0 for *Gallus gallus*^39^, HdrR for *Oryzias latipes*^40^) by using bowtie2 (version 2.2.6)^41^ with the following parameters ‘-k 4 -X 2000 --sensitive’. These genomes were downloaded from the Ensembl database (release 89)^42^. Picard (version 2.9.0; http://broadinstitute.github.io/picard/faq.html) was used to remove duplicate reads from among properly paired mapped reads, which had been filtered by using SAMtools (version 1.4)^43^. For the analysis with uniquely hit ATAC-seq reads, we extracted only uniquely aligned reads after duplicate reads had been removed. After read start positions were corrected to account for the 9-bp insert between the adaptors, which was introduced by Tn5 transposase^19^, the single 5’-most base of each reads was retained to obtain single-base resolution. Finally, for each sample, we randomly selected 20 million aligned reads to create a read-depth– controlled dataset for further analysis. For analyses without reads aligned to the mitochondrial genome and with uniquely hit ATAC-seq reads, we randomly selected 18 million aligned reads for further analysis.

### Identification of accessible chromatin regions

To identify accessible chromatin regions that were consistently present across three biological replicates, ATAC-seq peaks were called according to the method described by Daugherty *et al*^44^ by using MACS (version 2.1.1)^45^. At first, read data for three biological replicates were pooled into a single dataset. For these pooled data and those for each replicate, peak calling was performed by using MACS at the moderate threshold (--nomodel --extsize 50 --shift -25 -p 0.1 --keep-dup all). Of the peaks in the pooled data, we retained those that had at least 50% overlap with peaks in all three replicates.

### Whole-genome pairwise alignment

To estimate the evolutionary origin of each genomic region, we generated whole-genome pairwise alignments against different sets of animal genomes^8,38-40,46-70^ (Supplementary Table 2). LASTZ (version 1.04.00)^71^, axtChain, chainAntiRepeat, chainSort, chainPreNet, chainNet and netSyntenic^72^ were used for pairwise sequence alignments. The following parameters were used for LASTZ: ‘O=400 E=30 H=0 K=3000 L=3000 H=2200 T=1’. For axtChain, the parameters were ‘-minScore=3000 -linearGap=medium’ or ‘- minScore=5000 -linearGap=loose’, depending on the phylogenetic distance between the two examined species (Supplementary Table 2). Default parameters were used for other programs.

### Estimation of evolutionary origins of accessible chromatin regions

We used the aligned genomes to determine the evolutionary categories of accessible chromatin regions; when an accessible chromatin region had at least two different evolutionary origins, each was treated as an independent sub-region. We prepared two datasets to ensure that the methods we used to estimate the evolutionary origins of genomic regions did not alter our conclusions. One dataset was defined by the divergence time of the largest monophyletic group that shared the regions with similar sequences. This dataset did not include accessible chromatin regions that were lost in some lineages or species and were not shared by any monophyletic groups. In the other dataset, the evolutionary origins were defined by the most evolutionarily distant species that had the regions with similar sequences according to Dollo parsimony^73,74^.

### Cross-species transcriptome comparison

Previously published^9,75^ RNA-seq datasets were used to calculate early-to-late whole-embryonic gene expression profiles (DDBJ accession DRA003460 for mouse [*M. musculus*], chicken [*G. gallus*], Chinese softshell turtle [*P. sinensis*], western clawed frog [*X. tropicalis*] and zebrafish [*D. rerio*]; DRA005309 for medaka [*O. latipes*]). In brief, RNA-seq reads were aligned to each reference genome^8,38-40,51,70^ by using Tophat2 (version 2.0.14)^76^ with the following parameters: ‘-g 1 -N 3 -- read-edit-dist 3’. For paired-end RNA-seq data of medaka embryos, among all aligned reads, only properly paired mapped reads with a primary alignment were then filtered by using SAMtools (version 1.4)^43^. To obtain the expression levels of genes on the basis of these mapped datasets, FPKM (fragments per kilobase of transcripts per million fragments mapped) values were calculated by using Cufflinks (version 2.2.1)^77^ and a gene set retrieved from the Ensembl database^42^. For inter-species comparisons of orthologous gene expression levels, we used 1:1 orthologue information between each pair of species that was obtained from the Ensembl Compara Database through BioMart^78^. To evaluate transcriptome similarity between samples, a Spearman correlation coefficient was calculated by using the expression values of orthologous genes, as described by Wang *et al*^8^. As indicated previously^9, 79^, phylogenetic relationships between the species being compared were considered in the transcriptome-based identification of vertebrate-conserved stages. For stage combinations among the six different vertebrate embryos (mouse, chicken, softshell turtle, western clawed flog, zebrafish and medaka), we extracted pairs of species that reflected the phylogenetic scale of interest (i.e., vertebrates) and took their average value as expDist. To estimate the most-conserved stages, we first identified stage combinations with the most similar expression profiles of 1:1 orthologues (lowest 1% expDists) from all combinations. We then visualized the contribution of each stage in medaka embryogenesis to this top 1% of similarly staged embryo combinations as P_top_ (percentage of stage included in the top similar 1% of stage–embryo combinations). This calculation process was performed 100 times with randomly selected biological replicate data to derive statistically robust conclusions. When 1:1 orthologues were used in this analysis, genes that lacked 1:1 orthologue counterparts in any of the species (for example, because of gene loss in any of the species being compared) were ignored, as previously described^9^. The other dataset for calculating expDists was the orthologue-group-based gene expression profiles, in which expression levels of in-paralogs defined by orthoMCL^80^ were summed and further compared between species to calculate expression distances.

### Estimation of evolutionary origins of protein-coding genes

We used peptide sequences from mouse, chicken and medaka to estimate the evolutionary origins of protein-coding genes. After removing entries shorter than 30 amino acids, we selected the longest peptide sequence for each gene and used it in tblastn searches (version 2.7.1) with a threshold E-value < 10^−10^ to identify regions with similar sequences in the other species’ genomes. We then used this information to estimate the evolutionary origin of each protein-coding region according to the divergence time of the highest monophyletic group in which all examined species shared the regions with similar sequences in their genomes. At each developmental stage, the total expression level of genes with the same evolutionary age were calculated by using the previously determined FPKM-normalized whole embryonic gene expression profiles of mouse, chicken and medaka^9,75^. As with the estimation of the evolutionary origins of accessible chromatin regions, we prepared another dataset to ensure that the methods we used to estimate the evolutionary origins did not influence our conclusions. In this alternative dataset, the evolutionary origins of protein-coding genes were estimated according to the most evolutionarily distant species that had the regions with similar sequences in its genome, as we did earlier.

### Statistics

We used R (ver. 3.4.4)^81^ to perform all statistical analysis. An alpha level of 0.05 was adopted for statistical significance throughout the analyses. For the statistical analyses presented in Figure 2 and Extended Data Figures 5–10, the Kruskal–Wallis rank sum test was used. For the analysis presented in Extended Data Figures 1a–e and 3b, correlation coefficients were regarded as valid only when the comparison was confirmed to have a significant correlation by a test of no correlation. For statistical analyses presented in Extended Data Figure 1f, the Friedman rank sum test was used. Detailed information on statistical analyses used in this study is provided in Supplementary Tables 3–11.

### Software

In addition to software specified elsewhere in the Methods section, we used bedtools (version 2.27.1)^82^, ggplot2^83^, and Integrative Genomics Viewer^84^.

### Code availability

All analyses were done with publicly available software. Customised Ruby scripts for data analysis are available from the corresponding author upon request.

## Data availability

All ATAC-seq data generated during this study have been deposited in the DNA Data Bank of Japan and are accessible through accession number DRA006971.

## Extended Data

**Extended Data Figure 1.**
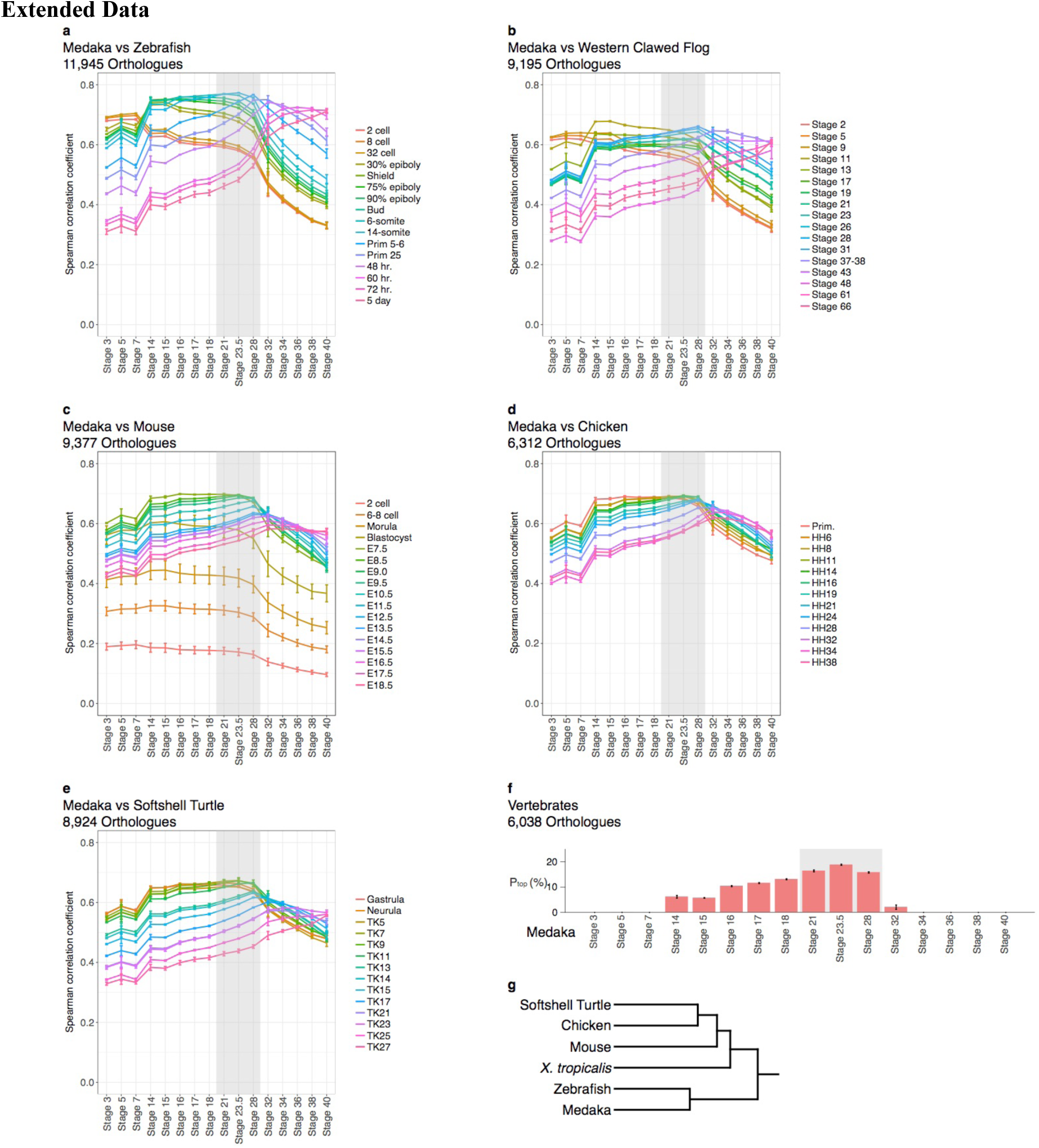
Conserved mid-developmental period during medaka embryogenesis identified by transcriptome similarity. **a–e.** All-to-all-stage comparisons of the transcriptome data of medaka embryos against zebrafish (**a**), western clawed flog (**b**), softshell turtle (**c**), chicken (**d**) and mouse (**e**) embryos are shown. The Spearman correlation coefficient was calculated by using FPKM values of orthologous genes to evaluate the transcriptomic similarity between samples. Lines were coloured according to the developmental stages of each species. The number of orthologous genes used for comparing transcriptomes between species is shown at the top of each panel. For all line graphs, data are presented as means ± 1 standard deviation of all pairs of biological replicates. **f.** The percentages of medaka developmental stages included in the most similar (lowest 1% of expDists) combinations of staged embryos from six vertebrates (mouse, chicken, softshell turtle, western clawed flog, zebrafish and medaka) are shown as P_top_ (see Methods for more details). Higher P_top_ values indicate developmental stages that are more highly conserved among the embryos of these vertebrates. expDist values were calculated to assess evolutionary conservation throughout the evolution of each of the six vertebrate species (see Supplementary Note 1 and Methods for more detail). To calculate expDists in the vertebrates, whole embryonic expression levels of 6,038 1:1 orthologues in the embryos of the six vertebrates were used^9,75^. Error bars represent the standard deviations for P_top_ values in 100 randomly selected biological replicates (BRI-exp). Changes in the P_top_ values of the developmental stages were statistically significant (Friedman test with 100 randomly selected BRI-exp for each species). **g.** Phylogenetic relationships of the six vertebrate species referenced in calculating expDists. Note that the developmental stages close to stage 24 of the medaka embryogenesis (highlighted in grey) were highly conserved across the examined vertebrate species. Sample sizes and detailed information regarding the statistical tests (test statistic, degrees of freedom and *P* value) are in Supplementary Table 4.

**Extended Data Figure 2.**
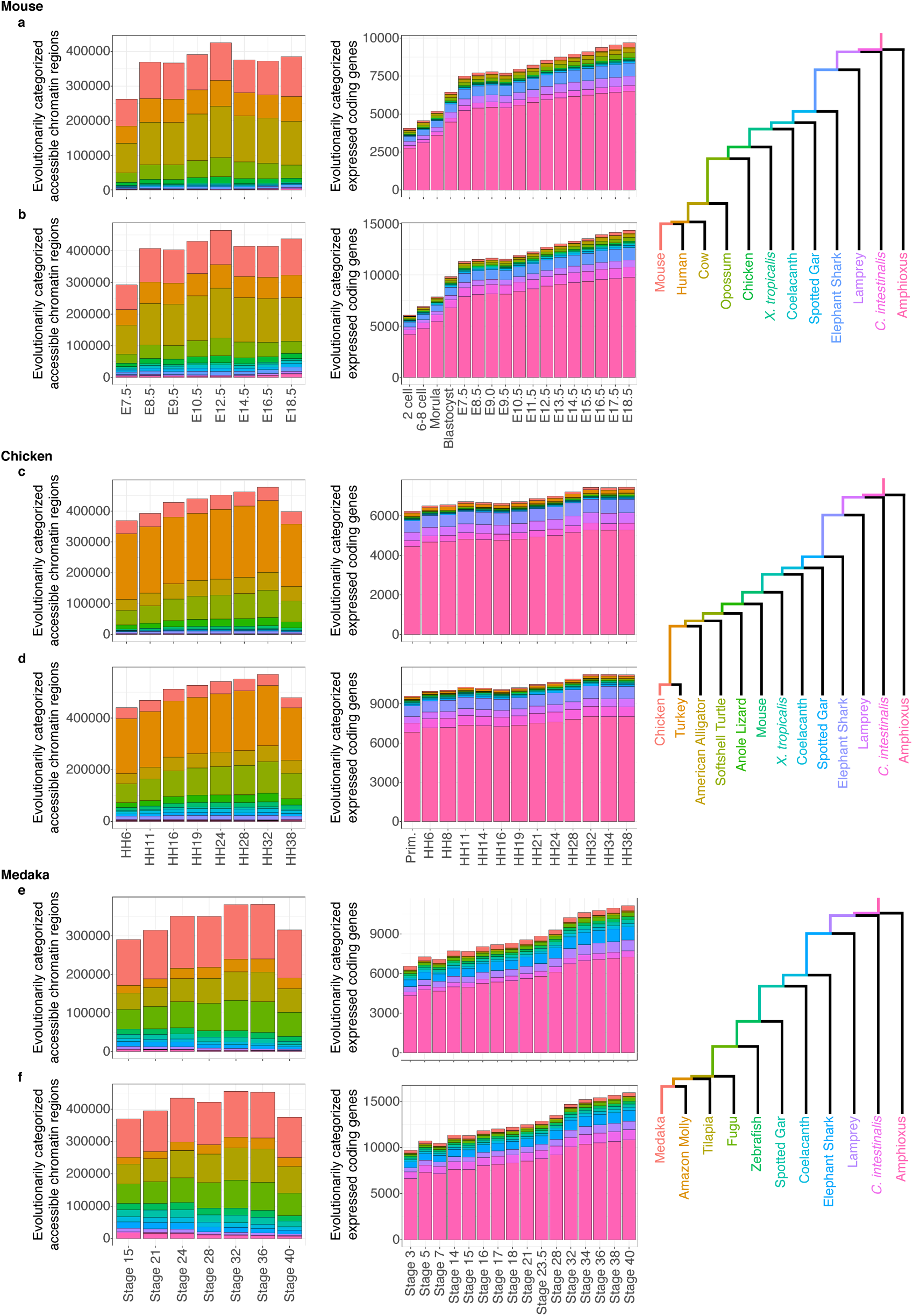
Numbers of accessible chromatin regions and expressed protein-coding genes categorized according to evolutionary origin. Stacked bar graphs show the numbers of evolutionarily categorized accessible chromatin regions (left) and expressed protein-coding genes (FPKM > 1; middle) for different developmental stages in mouse (**a, b**), chicken (**c, d**) and medaka (**e, f**). The evolutionary origin of genomic elements were estimated according to the largest monophyletic group that shared the regions with similar sequences (**a, c, e**) and according to the most evolutionarily distant species with the regions with similar sequences in its genome (**b, d, f**) (see Methods). Colours in each stacked bar graph represent the estimated evolutionary origin when each element emerged, and the colour corresponds to the tracks of the evolutionary trajectory, which are shown as a tree on the right. Phylogenetic relationships and split time were adopted from the TimeTree database^29^. Note that almost all regions with accessible chromatin in embryogenesis have been acquired since the origin of vertebrates, whereas the majority of expressed protein-coding genes were acquired before that.

**Extended Data Figure 3.**
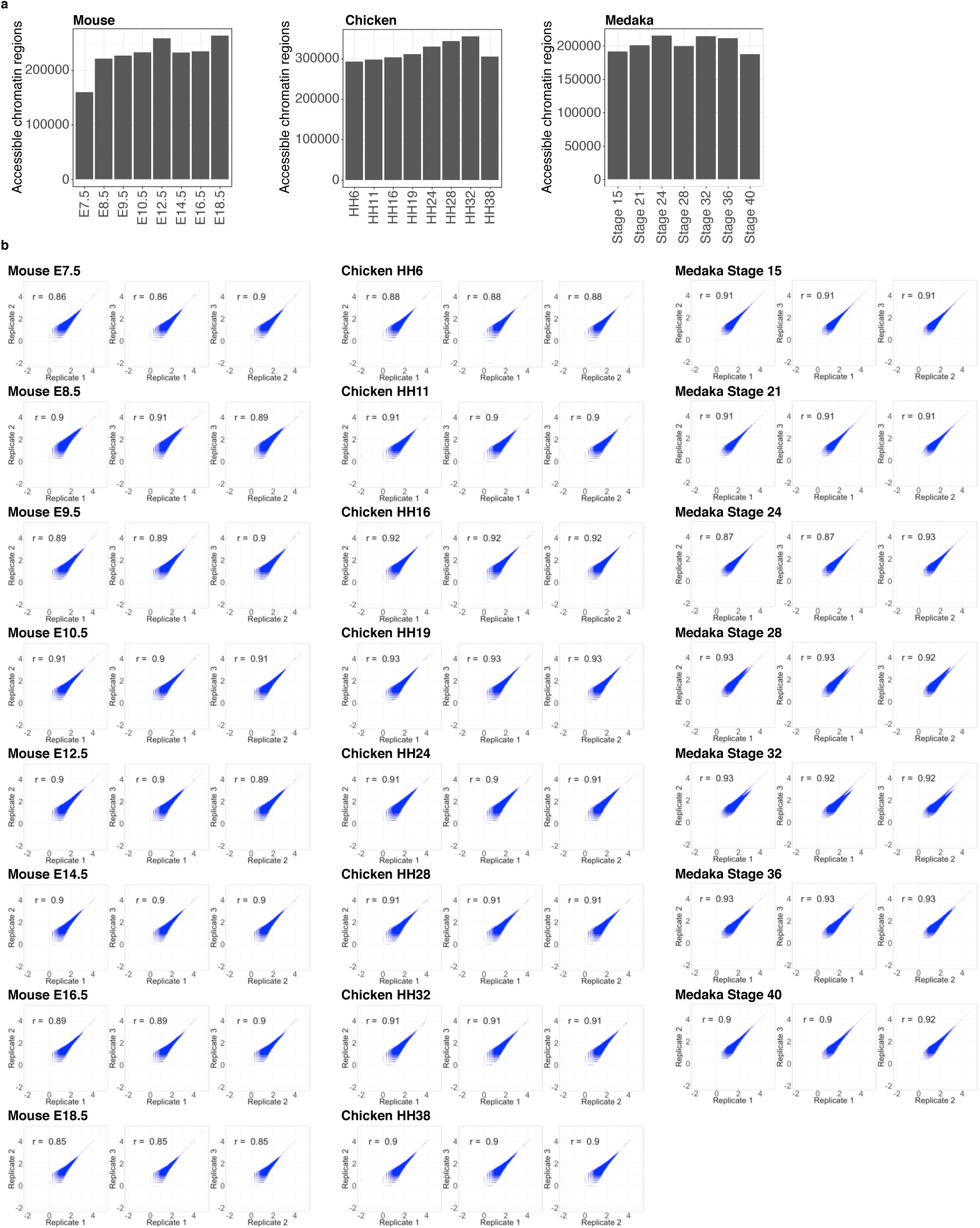
Numbers of accessible chromatin regions and correlations between biological replicates of whole-embryo ATAC-seq signals. **a.** Bar graphs show the numbers of accessible chromatin regions at different developmental stages. **b.** These scatter plots show correlations between the ATAC-seq signals (log_10_-RPM; reads per million mapped reads) at accessible chromatin regions for each developmental stage across three biological replicates. The Pearson correlation coefficient (r) is shown in each graph. Sample sizes and detailed information for the statistical tests (test statistic, degrees of freedom and *P* value) are shown in Supplementary Table 5.

**Extended Data Figure 4.**
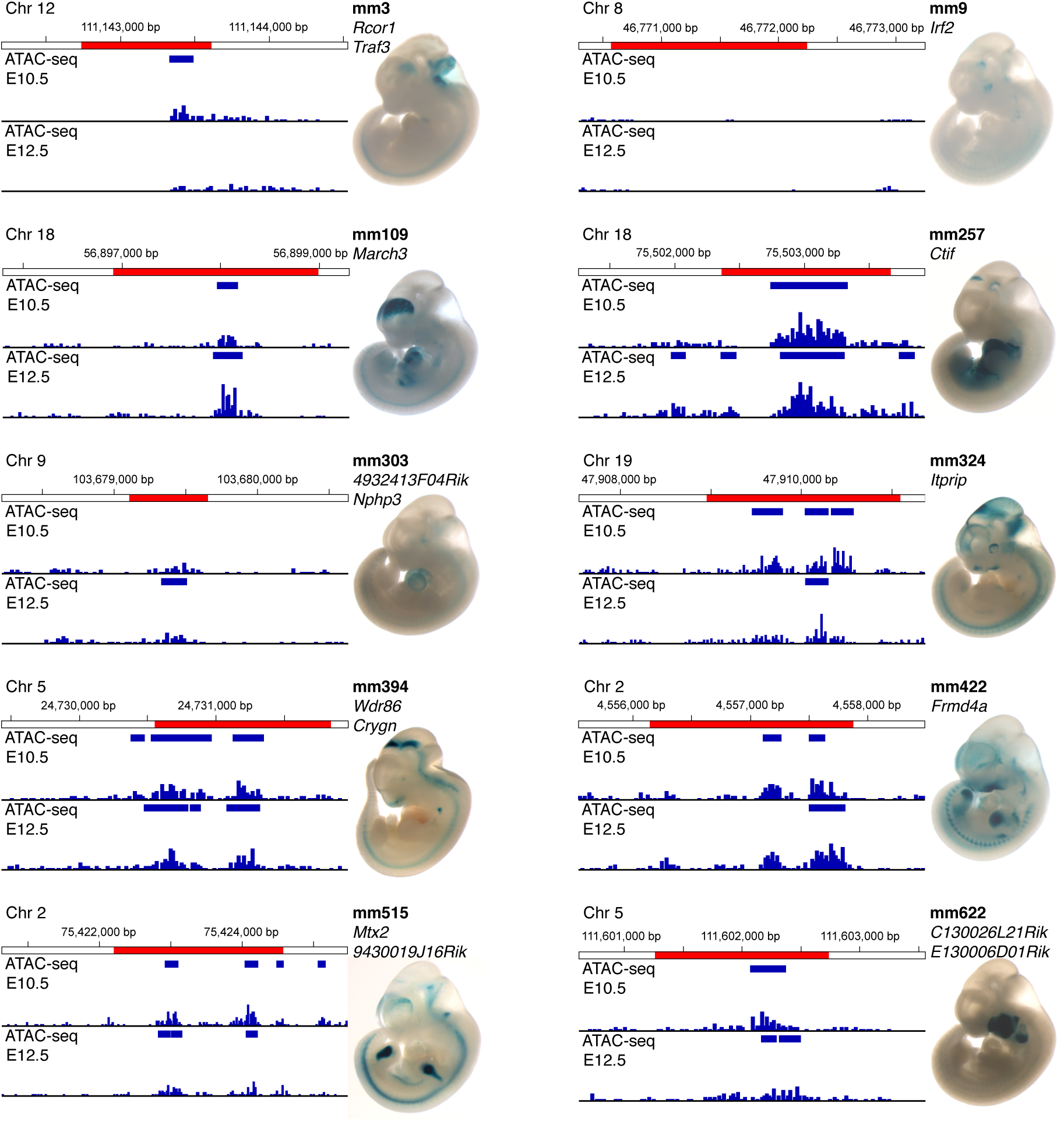
Enhancers that drive wider expression tend to have higher ATAC-seq signals. We selected 10 representative enhancers (red regions) from the VISTA Enhancer Database^30^ and checked for correlations between the whole-embryo ATAC-seq signal intensities and *in vivo* enhancer activity (*LacZ* reporter assay) in E11.5 mice. Images of embryos are from VISTA Enhancer Database^30^. For each annotated enhancer, the VISTA enhancer ID and the flanking gene are shown above the embryo image. The ATAC-seq signal shown in each panel represents the mean value of the three biological replicates. Blue boxes indicate the accessible chromatin regions identified by enrichment of the whole-embryo ATAC-seq signal across three biological replicates (see Methods section).

**Extended Data Figure 5.**
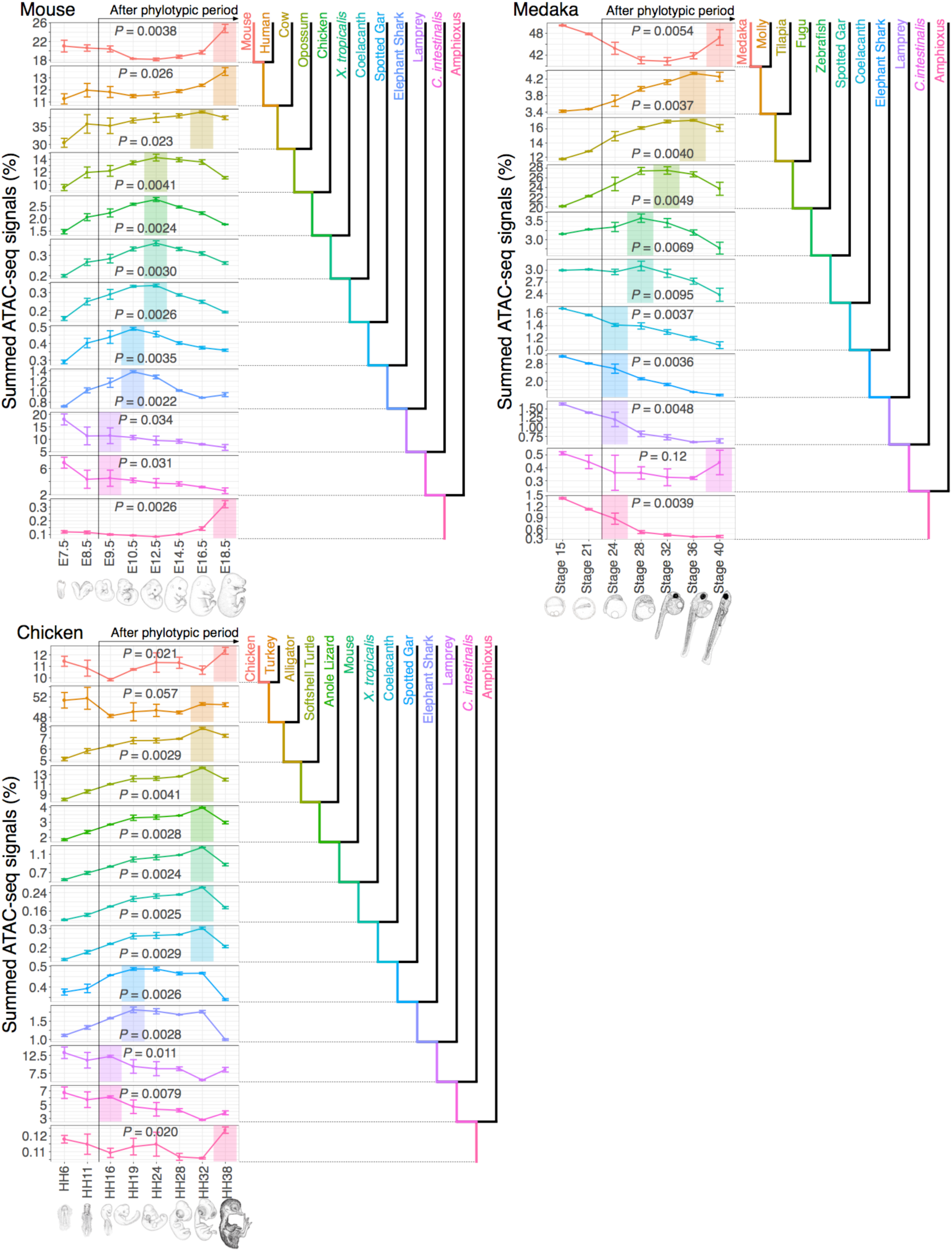
Transition of signal peaks in evolutionarily categorized chromatin accessibilities calculated by using uniquely hit ATAC-seq reads. The graphs presented were generated by using the same method as for Figure 2, except that uniquely hit ATAC-seq reads were used (see Methods). For each developmental stage in three vertebrate species (mouse, chicken, and medaka), the percentages corresponding to the summed signal intensity for each evolutionary category divided by the total signal intensity for all categories are shown. The colour of each category indicates the period of the evolutionary origin of the region, represented in the phylogenetic trees adjacent to the graphs. In each graph, developmental stages with the highest values after the potential phylotypic period are highlighted in the corresponding colour. Error bars indicate the standard deviations of three biological replicates for each developmental stage. The *P* value shown in each graph was calculated by using the Kruskal–Wallis rank sum test. The results shown in this figure are essentially the same as those shown in Figure 2. Sample sizes and detailed information regarding the statistical tests (test statistic, degrees of freedom and *P* value) are shown in Supplementary Table 6.

**Extended Data Figure 6.**
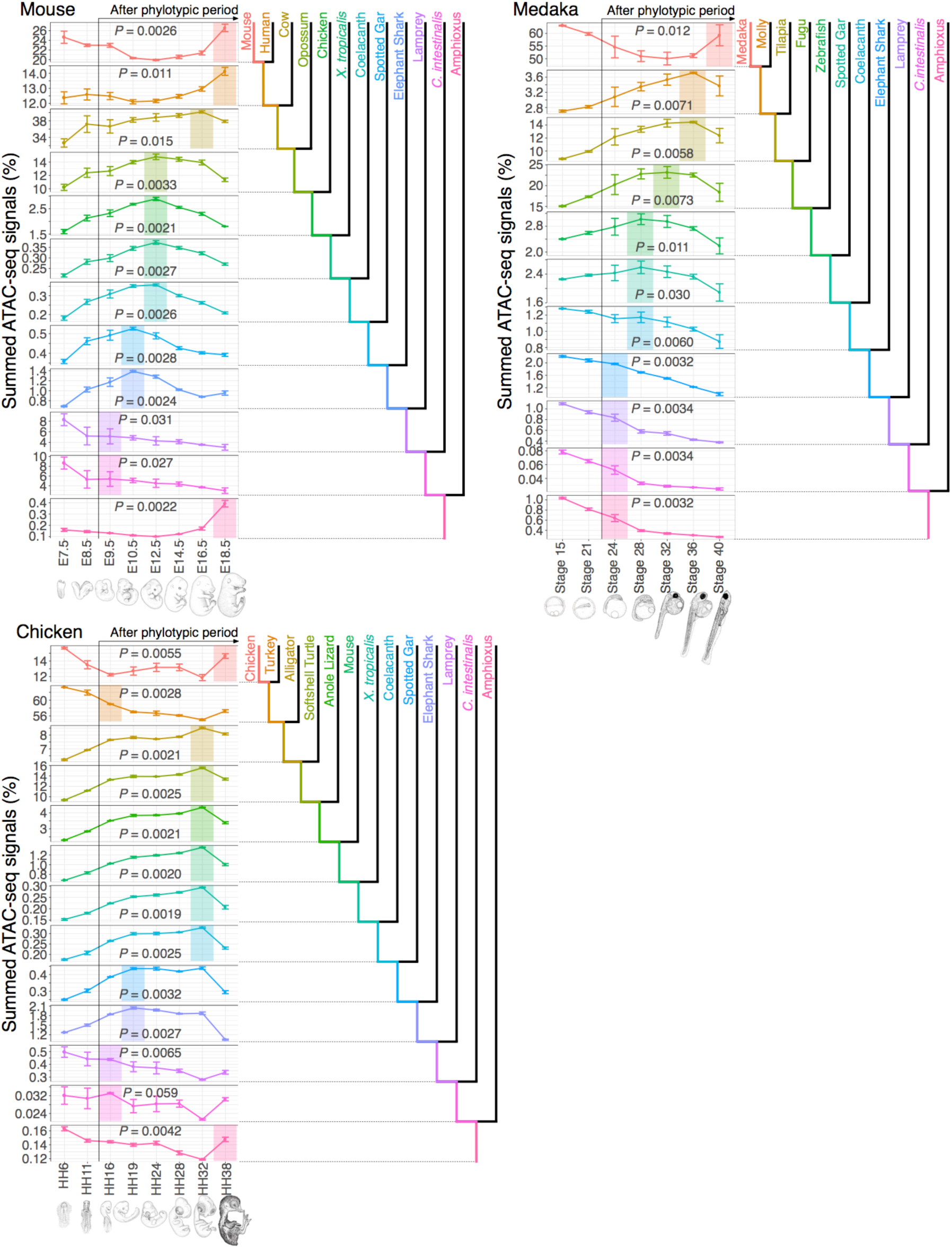
Transition of signal peaks in evolutionarily categorized chromatin accessibilities throughout the genome, except for the mitochondrial genome. The graphs presented were generated by using the same method as for Figure 2, except that ATAC-seq reads aligned to the mitochondrial genome were removed (see Methods). For each developmental stage in three vertebrate species (mouse, chicken, and medaka), the percentages corresponding to the summed signal intensity for each evolutionary category divided by the total signal intensity for all categories are shown. The colour of each category indicates the period of the evolutionary origin of the region, represented in the phylogenetic trees adjacent to the graphs. In each graph, developmental stages with the highest value after the potential phylotypic period are highlighted in the corresponding colour. Error bars indicate the standard deviation of three biological replicates for each developmental stage. The *P* value shown in each graph was calculated by using the Kruskal–Wallis rank sum test. The results shown in this figure are essentially the same as those shown in Figure 2 of the main data. Sample sizes and detailed information regarding the statistical tests (test statistic, degrees of freedom and *P* value) are shown in Supplementary Table 7.

**Extended Data Figure 7.**
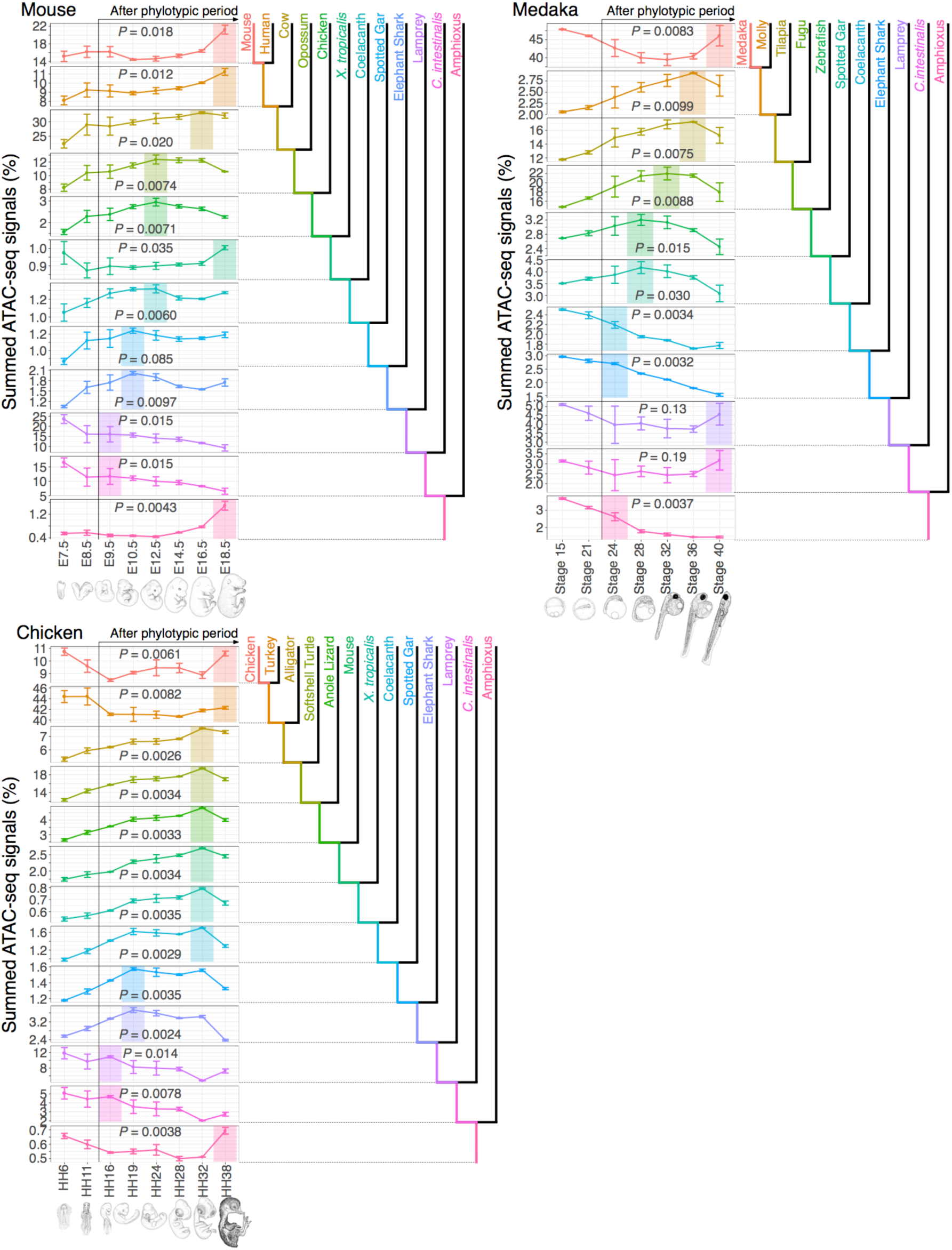
Transition of signal peaks in chromatin accessibilities categorized according to evolutionary origins based on Dollo parsimony. The graphs presented were generated by using the same method as for Figure 2, except for the way that evolutionary origins were estimated (see Methods). In short, the evolutionary origins of the examined regions were estimated according to the most evolutionarily distant species with aligned regions in its genome. For each developmental stage in three vertebrate species (mouse, chicken, and medaka), the percentages corresponding to the summed signal intensity for each evolutionary category divided by the signal intensity for all categories are shown. The colour of each category indicates the period of the evolutionary origin of the region, represented in the phylogenetic trees adjacent to the graphs. In each graph, developmental stages with the highest value after the potential phylotypic period are highlighted in the corresponding colour. Error bars indicate the standard deviation of three biological replicates for each developmental stage. The *P* value shown in each graph was calculated by using the Kruskal–Wallis rank sum test. Sample sizes and detailed information regarding the statistical tests (test statistic, degrees of freedom and *P* value) are shown in Supplementary Table 8.

**Extended Data Figure 8.**
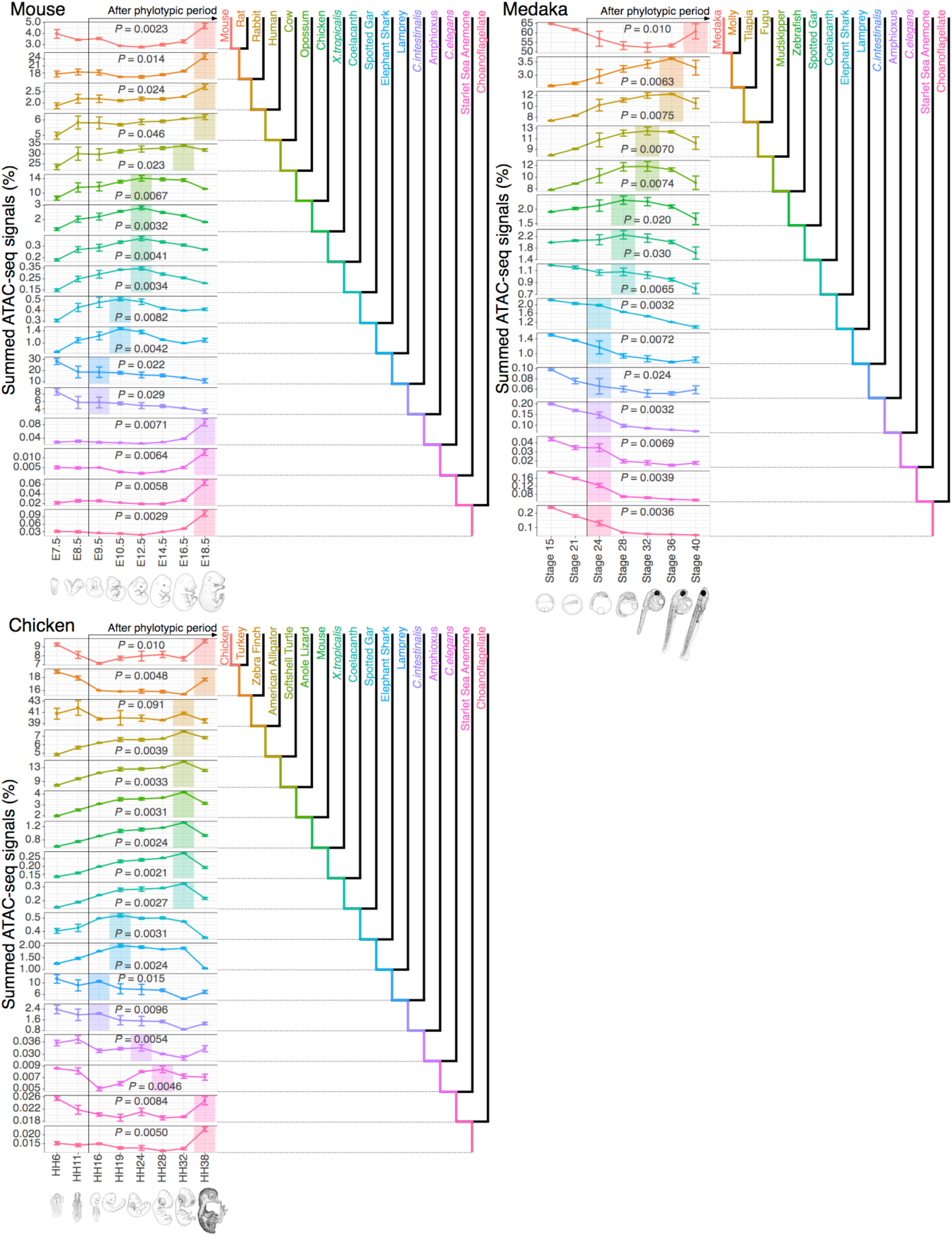
Transition of signal peaks in chromatin accessibilities categorized according to evolutionary origins estimated by using different genome sets. The graphs presented were generated by using the same method as for Figure 2, except with different sets of genomes. For each developmental stage in three vertebrate species (mouse, chicken, and medaka), the percentages corresponding to the summed signal intensity for each evolutionary category divided by the signal intensity for all categories are shown. The colour of each category indicates the period of the evolutionary origin of the region, represented in the phylogenetic trees adjacent to the graphs. In each graph, developmental stages with the highest value after the potential phylotypic period are highlighted in the corresponding colour. Error bars indicate the standard deviation of three biological replicates for each developmental stage. The *P* value shown in each graph was calculated by using the Kruskal–Wallis rank sum test. Sample sizes and detailed information regarding the statistical tests (test statistic, degrees of freedom and *P* value) are shown in Supplementary Table 9.

**Extended Data Figure 9.**
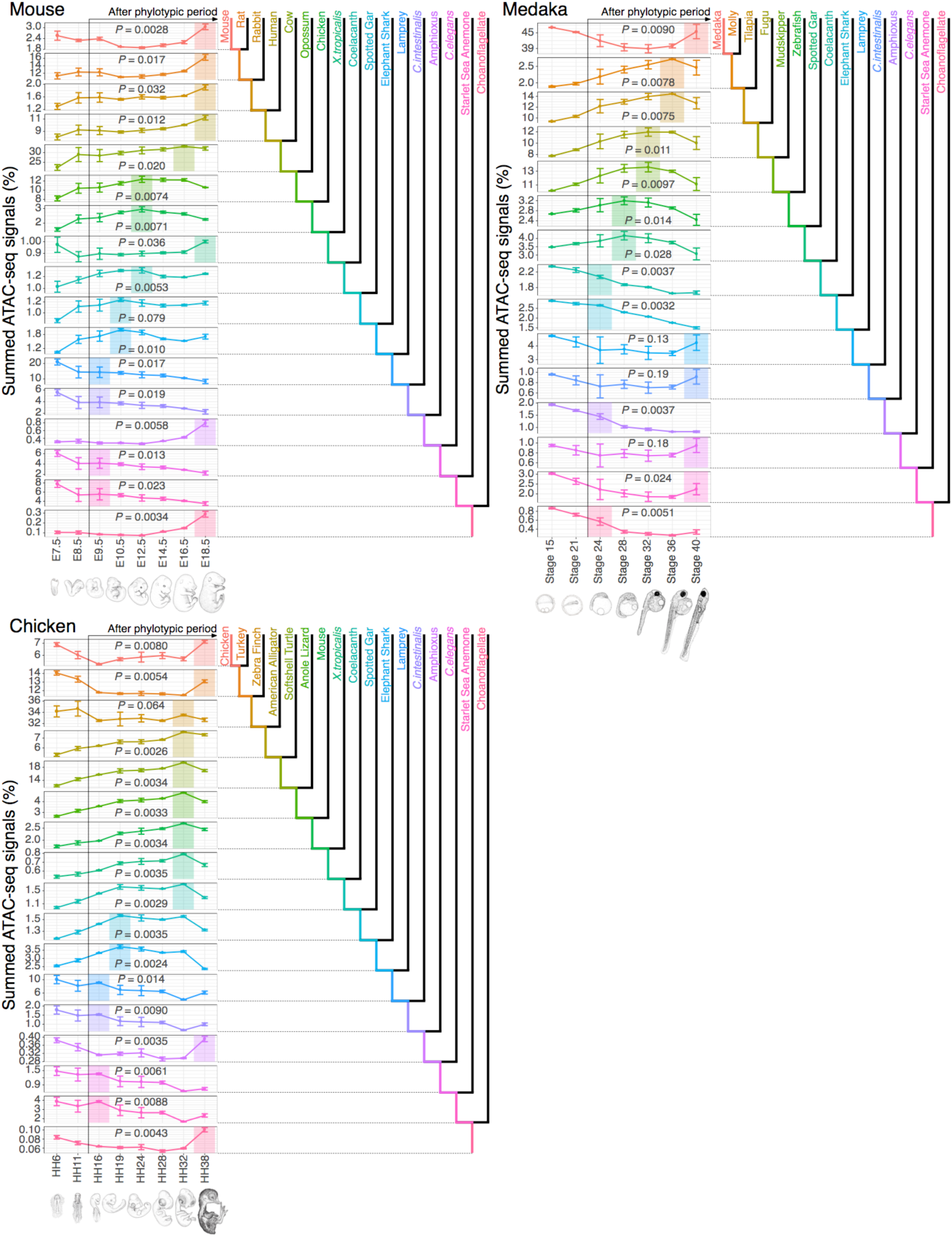
Transition of signal peaks in chromatin accessibilities categorized according to evolutionary origins estimated by using Dollo parsimony and different genome sets. The graphs presented were generated by using the same method as for Extended Data Figure 8, except for the way that evolutionary origins were estimated (see Methods). In short, the evolutionary origins of the examined regions were estimated according to the most evolutionarily distant species with the regions with similar sequences in its genome. For each developmental stage in three vertebrate species (mouse, chicken, and medaka), the percentages corresponding to the summed signal intensity for each evolutionary category divided by the signal intensity for all categories are shown. The colour of each category indicates the period of the evolutionary origin of the region, represented in the phylogenetic trees adjacent to the graphs. In each graph, developmental stages with the highest value after the potential phylotypic period are highlighted in the corresponding colour. Error bars indicate the standard deviation of three biological replicates for each developmental stage. The *P* value shown in each graph was calculated by using the Kruskal–Wallis rank sum test. Sample sizes and detailed information regarding the statistical tests (test statistic, degrees of freedom and *P* value) are shown in Supplementary Table 10.

**Extended Data Figure 10.**
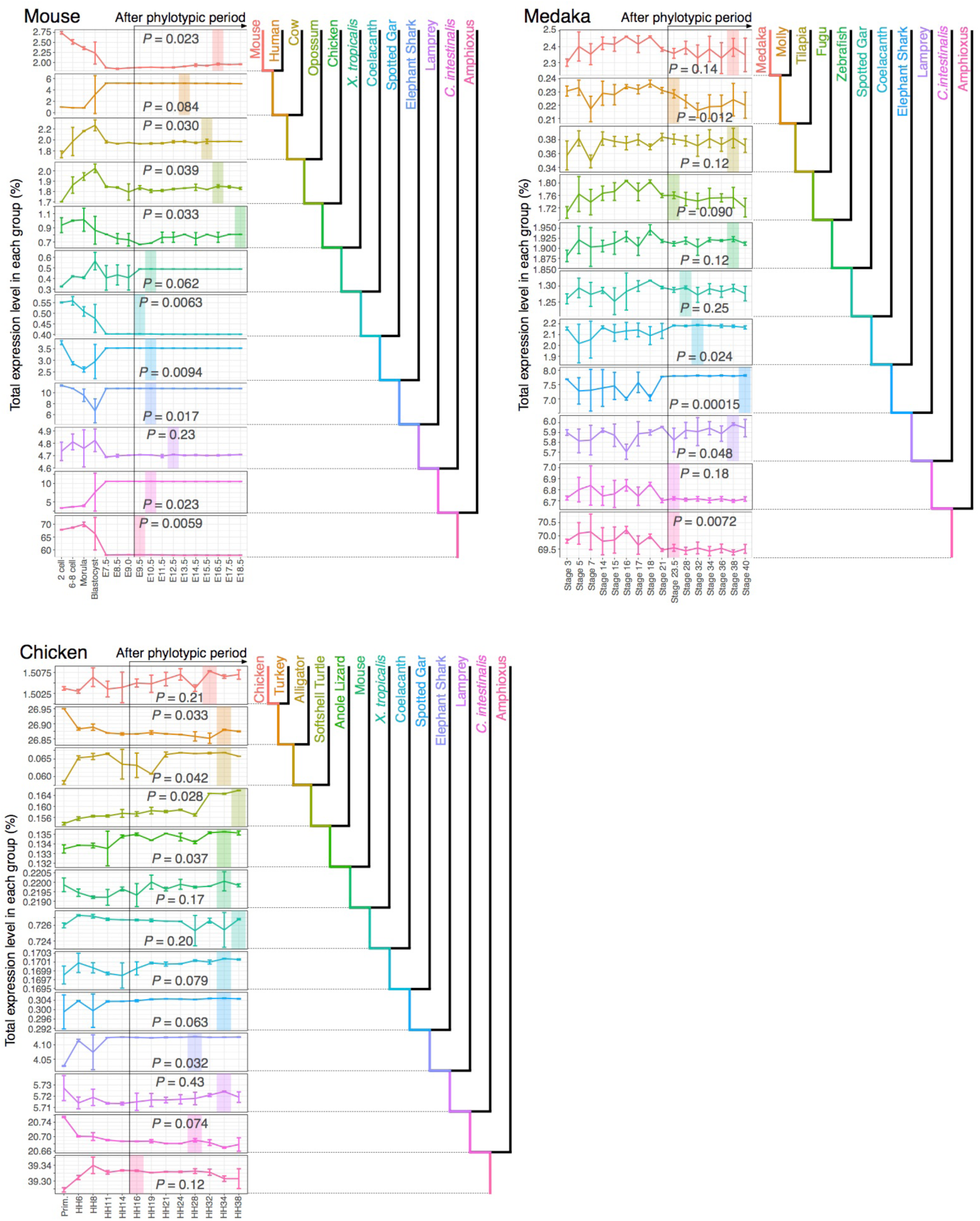
Transition of signal peaks in total expression levels of evolutionarily categorized protein-coding genes. For each developmental stage in three vertebrate species (mouse, chicken, and medaka), the percentages corresponding to the summed expression level for each evolutionary category divided the total expression level for all categories are shown. The colour of each category indicates the period of the evolutionary origin of the regions, represented in the phylogenetic trees adjacent to the graphs. In each graph, developmental stages with the highest value after the potential phylotypic period are highlighted in the corresponding colour. Error bars indicate the standard deviation of biological replicates for each developmental stage. The *P* value shown in each graph was calculated by using the Kruskal–Wallis rank sum test. Sample sizes and detailed information regarding the statistical tests (test statistic, degrees of freedom and *P* value) are shown in Supplementary Table 11.

