## Supplementary material for "Evolutionary transition in accessible chromatin landscapes during vertebrate embryogenesis"

### Supplementary Note 1
Previous studies identified stages around HH16 in chicken and E9.0 in mouse as the most transcriptomically conserved periods among vertebrates[6,8,9] and proposed these stages as the potential phylotypic period in these species. Because no such stages had been reported for medaka, we performed a cross-species transcriptome comparison, using a previously described method[6,8,9]. In this analysis, we used early-to-late embryonic transcriptome data collected for six vertebrate species—mouse (*M. musculus*), chicken (*G. gallus*), softshell turtle (*Pelodiscus sinensis*), western clawed frog (*Xenopus tropicalis*), zebrafish (*Danio rerio*) and medaka (*O. latipes*)—in previous studies[8,9]. Pairwise comparisons of the gene expression profiles of 1:1 orthologues showed that medaka at around stage 24 tends to have increased expression similarity to other vertebrate embryos (Extended Data Figure 1a–e). In addition, the stage 24 we identified here matches the stage that shows the most similar anatomical pattern between medaka and zebrafish and has been proposed as a potential phylotypic period in medaka[85] (around the pharyngula stage). By calculating expression distances[9] (expDists; see Methods) with 1:1 orthologues (6038 orthologues in total) taking phylogeny into consideration, we further demonstrated that the stages around stage 24 are the period showing the highest conservation across vertebrate species in medaka embryogenesis. In short, among the most similar (lowest 1% of expDists) combinations of stages, stage 24 was the developmental stage most frequently observed in medaka (Extended Data Figure 1f–g). Meanwhile, using 1:1 orthologues for the comparison might introduce bias, as previously acknowledged[9], because several orthologue counterparts were removed. We therefore used orthologue groups to calculate expDists, to cover all genes in each species and thus remove bias in gene repertoire. In brief, the method assigns a '0' expression level to the genes of species in which the genes have no orthologous counterparts and sums the expression levels of genes within the same orthologue group (in-paralogs). Using this orthologue group-based method, we obtained essentially the same result as one using 1:1 orthologues (data not shown). Taken together, these results suggest that medaka embryogenesis shows hourglass-like divergence and that the period around stage 24 is the potential phylotypic period in this species.

### Supplementary Note 2
To test whether the recapitulative tendency observed in Figure 2 was corroborated by analyses with different parameters, we prepared four additional datasets, as follows.

***2.1 Uniquely hit ATAC-seq reads (Extended Data Figure 5)***

Because some ATAC-seq reads mapped to multiple genomic regions (e.g., transposable elements,[86] which have many copies in the genome), we used the best-hit genomic region for the analysis in Figure 2. To test whether this best-hit parameter biased the result, we prepared another dataset made of uniquely hit ATAC-seq reads, which excluded ATAC-seq reads that were aligned to multiple regions in the genome. This dataset yielded essentially the same results as in Figure 2 (Extended Data Figure 5).

***2.2 Without ATAC-seq reads mapped to the mitochondrial genome (Extended Data Figure 6)***

ATAC-seq reads can contain various proportions of sequences from mitochondrial genomes[34]. In our ATAC-seq samples, the average mitochondrial fractions were 8.7%, 7.7% and 2.9% of properly mapped reads (duplicate reads not included) in mouse, chicken and medaka embryos, respectively. Because these percentages were low compared with those in other studies using ATAC-seq without any treatment for mitochondrial DNA depletion (between 10% and >50%[34]; between 30% and 50%[87]; >80%[88]; between 20% and 80%[89]), we did not exclude ATAC-seq reads mapped to the mitochondrial genome to perform the analysis shown in Figure 2. However, we cannot exclude the possibility that including these fractions influenced the observed recapitulative tendency. We therefore removed ATAC-seq reads that were mapped to the mitochondrial genome and assessed whether the recapitulative tendency was still apparent. For chicken, although the developmental stage with the signal peak in the second-newest category changed from HH38 to HH16 (Extended Data Figure 6), the overall tendency was essentially the same as that in Figure 2. For medaka, although including mitochondrial sequences altered the older limit of evolutionary range over the recapitulative tendency during late embryogenesis (Extended Data Figure 6), the same tendency was observed for regions at least newer than the vertebrate–urochordate split.

***2.3 With evolutionary origins based on Dollo parsimony (Extended Data Figure 7)***

In Figure 2, the evolutionary categories of accessible chromatin regions were classified according to the divergence time of the largest monophyletic group that shared the regions with similar sequences. However, this criterion overlooks regions that were lost in some lineages or species. Similarly, the inclusion of low-quality genomic sequence data could affect the evolutionary categories for each region. We therefore prepared another dataset to test whether applying different criteria for the evolutionary categorization would affect the observed tendency. In this dataset, the evolutionary origins of accessible chromatin regions were estimated according to the most evolutionarily distant species that had the regions with similar sequences based on Dollo parsimony[73,74], which allows multiple losses but only one gain. Although the signal peak of coelacanth–*X. tropicalis* in mouse embryogenesis did not follow the recapitulative tendency, the overall tendency observed from this dataset (Extended Data Figure 7) was essentially the same as those shown in Figure 2.

***2.4 Genome sets including more evolutionarily distant ones for estimating evolutionary origins of genomic regions (Extended Data Figures 8, 9)***

To study the evolutionary acquisition of regulatory regions during vertebrate evolution, we selected chordate species genomes to estimate the evolutionary origins of genomic regions (Figure 2). However, whether the recapitulative tendency would be similar to that in Figure 2 when more evolutionarily distant species were included was unclear. This analysis is also important for determining the evolutionarily older limit of the recapitulative tendency, especially for mouse and chicken embryogenesis. For these two species, using only the data shown in Figure 2 and Extended Data Figures 5–7, we cannot conclude that regions newer than the vertebrate–urochordate split show sequential transitions of chromatin accessibilities after recapitulation, because the second-oldest category of regions (the second from the bottom in each panel of Figure 2 and Extended Data Figures 5–7) contains both the regions for which the origins are between the divergence time of olfactores and that of vertebrates, but also older regions that have been lost from the cephalochordate lineage. In contrast as with medaka, we can conclude that the transitional pattern of chromatin accessibility of regions newer than the gnathostome–cyclostome split consistently shows a recapitulative tendency, because, in the analysis for Extended Data Figure 7, the signal peaks of the third-oldest category did not follow the tendency. Therefore, we performed the same analysis but with genomes of different apoikozoan (consisting the Animalia and the Choanoflagellatea) species. Here, we estimated the evolutionary origins of the accessible chromatin regions according to two different methods, as described in Supplementary Note 2.3 and the Methods section (Extended Data Figures 8, 9). For mouse and chicken embryogenesis, we noted the recapitulative tendency in chromatin accessibilities, similar to that shown in Figure 2, and found that the regions showing this tendency were newer than the vertebrate–urochordate split (Extended Data Figures 8, 9). In contrast, regarding medaka, although the same tendency was observed in later embryogenesis, the older limit of the evolutionary range for this tendency differed depending on the method used to estimate evolutionary origin, and on the genome set used for this analysis (Figure 2 and Extended Data Figures 8, 9). However, the chromatin accessibilities of regions at least newer than the cyclostome–gnathostome split consistently followed the recapitulative tendency, indicating that this tendency itself was robust against the genome set used.

### Supplementary Note 3

After teleosts diverged from other vertebrates, they experienced whole-genome duplication in the stem lineage[90]. Moreover, after the teleost genome duplication, eight major interchromosomal rearrangements took place in the stem-teleosts[40], and the genomic sequence evolved rapidly compared with the Holostei and Sarcopterygii[52-54,91]. These events might cause difficulties in estimating the origins of accessible chromatin regions—especially older ones—in the medaka genome. In addition, they might be part of the reason why the older limit of the evolutionary range for the transition during medaka late embryogenesis was affected by the genome set used, as well as by the methods used to estimate the evolutionary origins of regions (Figure 2 and Extended Data Figures 7–9).
